# Mental geometry of 3D size and shape perception

**DOI:** 10.1101/761650

**Authors:** Akihito Maruya, Qasim Zaidi

**Author notes:** **AUTHOR CONTRIBUTIONS:** AM and QZ designed the study. AM programmed and ran the experiments. AM & QZ analyzed and modeled the results. AM & QZ wrote the paper.

## Abstract

Judging poses, sizes and shapes of objects accurately is necessary for organisms and machines to operate successfully in the world. Retinal images of 3D objects are mapped by the rules of projective geometry, and preserve the invariants of that geometry. Since Plato, it has been debated whether geometry is innate to the human brain, and Poincare and Einstein thought it worth examining whether formal geometry arises from experience with the world. We examine if humans have learned to exploit projective geometry to estimate sizes and shapes of objects in 3D scenes.

Numerous studies have examined size invariance as a function of physical distance, which changes scale on the retina, but surprisingly, possible constancy or inconstancy of relative size seems not to have been investigated for object pose, which changes retinal image size differently along different axes. We show systematic underestimation of length for extents pointing towards or away from the observer, both for static objects and dynamically rotating objects. Observers do correct for projected shortening according to the optimal back-transform, obtained by inverting the projection function, but the correction is inadequate by a multiplicative factor. The clue is provided by the greater underestimation for longer objects, and the observation that they appear more slanted towards the observer. Adding a multiplicative factor for perceived slant in the back-transform model provides good fits to the corrections used by observers. We quantify the slant illusion with relative slant measurements, and use a dynamic demonstration to show the power of the slant illusion.

In biological and mechanical objects, distortions of shape are manifold, and changes in aspect ratio and relative limb sizes are functionally important. Our model shows that observers try to retain invariance of these aspects of shape to 3D rotation by correcting retinal image distortions due to perspective projection, but the corrections can fall short. We discuss how these results imply that humans have internalized particular aspects of projective geometry through evolution or learning, and how assuming that images are preserving the continuity, collinearity, and convergence invariances of projective geometry, supplements the Generic Viewpoint assumption, and simply explains other illusions, such as Ames’ Chair.

## Introduction

A biological or machine visual system can successfully operate in the world only by accurately judging poses, sizes and shapes of objects. In eyes with lenses, projective geometry describes the mapping of 3D scenes to retinal images, so in understanding what they are viewing, animals and humans have constant experience with the consequences of this geometry. Not surprisingly, whether geometrical operations are innately embedded in the human mind has been debated since antiquity, e.g. Plato’s Meno (Cooper, 2002). Poincare (1905) and Einstein (1921) expanded the debate to whether formal geometry arises from everyday experience. Consequently, showing that human performance in object size and shape estimation is based on geometric knowledge, could provide answers to age-old questions about the links between geometry and experience.

A 3D object seen from different views forms quite different retinal images, and many different objects can form identical retinal images (Ittelson, 1952), so 3D inferences based solely on monocular 2D retinal information are underspecified. However, the frequently occurring projection of reflections from objects on the ground to retinal images is a 2D-to-2D mapping, described by an invertible trigonometric function. So in this case, the back-transform derived by inverting the projection function could be used to make veridical inferences from retinal images. We showed previously that human observers consistently apply the optimal back-transform for pose inferences in 3-D scenes and in obliquely viewed pictures of 3D scenes (Koch et al 2018). This leads to veridical estimates for real 3D scenes, albeit with a systematic fronto-parallel bias, whereas pictured scenes are seen with an illusory rotation towards the observer. In the images we used for pose-estimation, we noticed that perceived sizes and shapes of objects appear distorted, especially those aspects of shape that depend on relative sizes in different directions. Here we tackle the mental geometry of estimating relative sizes using graphic displays calibrated against real objects. Variations in size estimates for different poses also provide information about perceived shape distortions, such as aspect ratios and relative sizes of limbs.

Shape is the geometrical attribute of an object that is invariant to translation, rotation, and scale effects (Kendal et al., 1999), but whether these invariances hold for perceptions of particular solid 3D shapes is an empirical question. Invariance to translation and rotation are properties of Euclidean spaces, whereas retinal image formation is described by perspective projection, which does not allow for these invariances. Perceptual invariance would thus require neural processes that overcome distortions created by the retinal projection, so the first step is to quantify perceived aspects of 3-D shape under different views.

Surprisingly, this issue has not been dealt with thoroughly, as almost all perceptual experiments have examined placing identical objects at different distances from the observer, which scales the size of the retinal image, but does not test for translation or rotation invariance. Gibson (1950) examined size invariance as a function of physical distance, and maintained that we can see approximately the true size by making use of inverse projection to recover the structure of the environment from the structure in the optic array. A number of studies have shown that humans are able to compensate partially for the retinal projection, so estimated sizes are closer to the physical size of the object (Gibson & Cornsweet, 1952; Joynson & Newson, 1962; Kaiser, 1967; Purdy, 1960; Sedgwick, 1983; Wallach & Moore, 1962). However, studies that asked observers to match a width to a depth found inconstancy of perceived 3D relative size, as depth had to be set 1.5 to 5 times the width to be judged as equal over different distances (Loomis, Da Silva, Fujita, & Fukushima, 1992, Beusmans,1998). In scene perception, a similar systematic perceptual anisotropy of depth versus width has been found (Baird & Biersdorf, 1967; Levin & Haber, 1993; Loomis & Philbeck, 1999; Norman et al, 1996; Philbeck, 2000; Ribeiro, Fukusima, & Da Silva, 1995; Toye, 1986; Wagner, 1985), suggesting a common cause, possibly insufficient correction of image compression caused by perspective projection. This perceptual anisotropy is not found when distances are estimated by non-visual motor tasks such as blind walking (Elliott, 1986, 1987; Fukusima, Loomis, & Da Silva, 1997; Loomis et al., 1992; Loomis, Klatzky, Philbeck, & Golledge, 1998; Philbeck & Loomis, 1997; Philbeck, Loomis & Beall, 1997; Rieser, Ashmead, Talor, & Youngquist, 1990; Sinai, Ooi, & He, 1998; Steenhuis & Goodale, 1988; Thomson, 1983), thus characterizing the anisotropy as related to the greater compression ratios for depths in retinal images as compared to widths.

Possible constancy or inconstancy of relative size seems not to have been investigated for object poses, which change retinal image size differently along different axes, so we address that lacuna in this study. The movie in Figure 1 shows a rigid 3D object with two physically equal arms at a right angle, rotating on the ground. To most people the arm pointing at or away from them appears transitorily shorter than the orthogonal arm. Using quantitative measurements, we show that for estimating relative sizes, observers’ generally correct for projective distortions according to the optimal back-transform, except for poses close to the line of sight. Size underestimation increases with object length, which we show is due to a slant illusion caused by projections of increased length being similar to increased slant, leading to possible overestimation of viewing elevation. We discuss how these results imply that humans have internalized particular aspects of projective geometry through evolution or learning.

**Figure 1.**
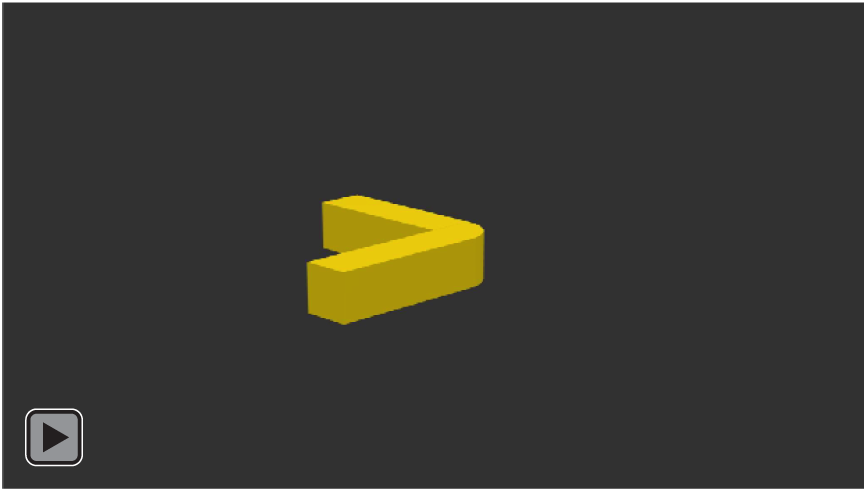
Demonstration of size inconstancy. A rigid object with two physically equal arms at a right angle is rotating on the ground. The arm pointing at or away from the observer appears transitorily shorter than the orthogonal arm.

### Experiment 1: Size estimates of 3D objects

The first question we address is how well observers can estimate 3D sizes across different poses of the same object lying on the ground, because projections lead to varying retinal image compression as a function of pose angle (Figure A1). Using Blender, we created a blue rectangular 3D parallelepiped (test stick) lying on the center of a dark ground, and a yellow vertical 3D cylinder (measuring stick) standing on the test stick. Blue parallelepipeds were presented in one of 16 poses from 0° to 360° every 22.5° (Figure 2a), with length equal to 10, 8 or 6 cm with 3×3 cm cross-section (Figure 2b). Observers estimated physical lengths by adjusting the height of the vertical cylinder between 2.75 and 12 cm, until the two appeared to be the same 3D length. In Experiment 1a the object was presented on a dark ground plane, and in Experiment 1b a regular white grid was superimposed on the ground plane to make it explicit (Figure 2c).

**Figure 2.**
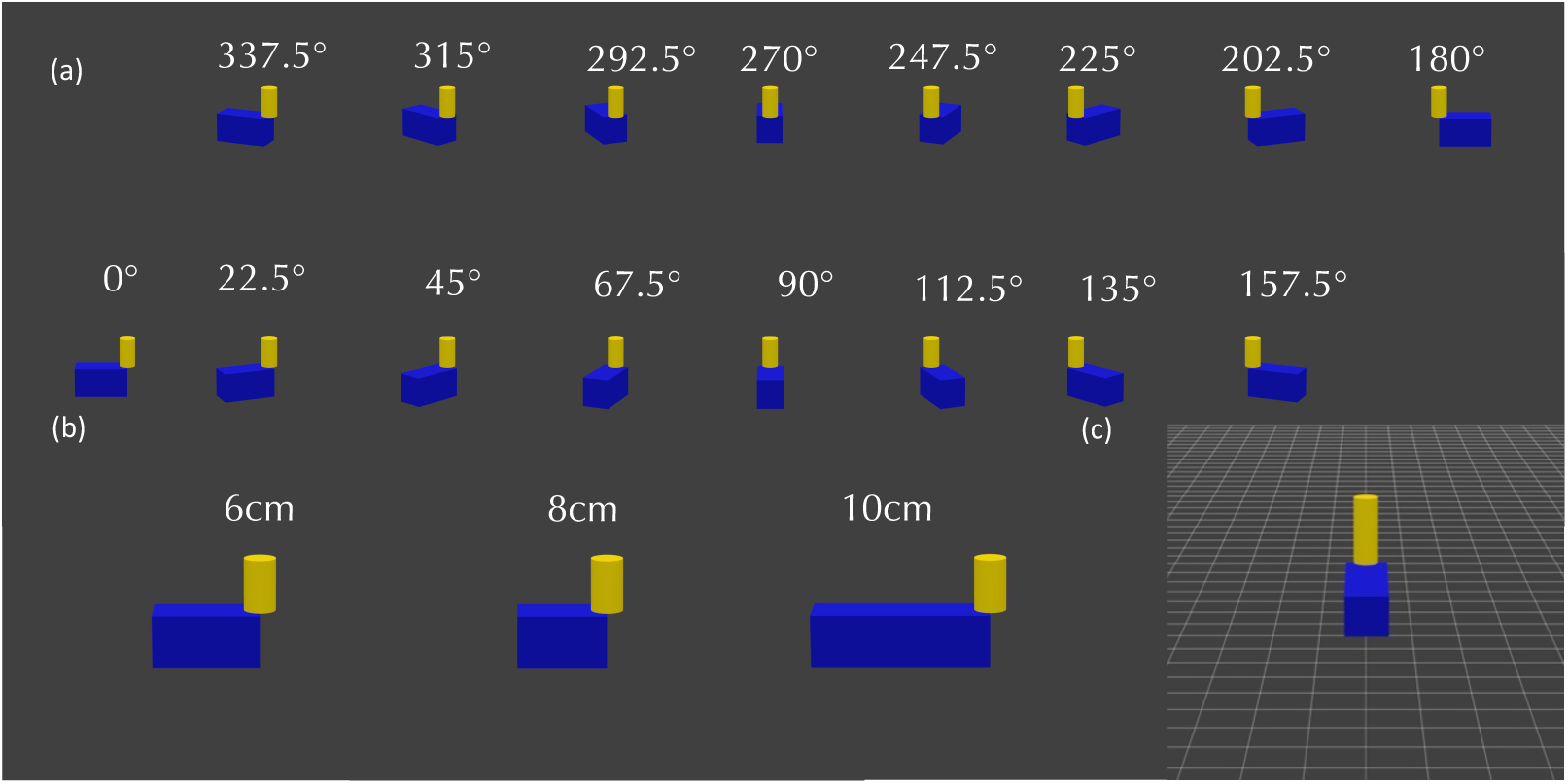
Stimuli for Experiment 1. (a) Experiment 1a: Blue test stick lying on dark ground in 16 different poses, from 0° to 337.5° at every 22.5°. (b) Experiment 1a: Three lengths, 6, 8, 10 cm, of blue stick were presented in each pose, with adjustable yellow measuring stick. (c) Experiment 1b: Same as Experiment 1a, except for a grid making the ground plane explicit.

#### Stimuli and Methods

The observer’s head was fixed in position by using a chinrest so that the image was viewed in front of the monitor, with an elevation angle of 15° at a distance of 1.0 m, matching the rendering parameters of the Camera in Blender. Displayed sizes in the Blender rendered images were calibrated against exact geometrical derivations to ensure accuracy of the simulations (Appendix). The retinal image was thus identical to that from the 3-D object, except for the absence of stereo disparity. Observers were instructed to equate the physical lengths of the two limbs by pushing buttons to adjust the height of the measuring stick. There were no time limits. Randomly ordered trials were repeated in three sets, and observers were allowed to take a break between sets. The line of sight through the center of the ground was designated the 90°-270° axis and the line orthogonal to it, the 0°-180° axis. Images were displayed on a 22 inch DELL SP2309W Display. Matlab and PsychoToolbox were used to display the stimulus, run the experiment, and analyze the data for all the experiments. 6 observers with normal or corrected vision participated. Viewing was binocular because Koch et al (2018) had not found any difference between monocular and binocular viewing for pose estimation in similar conditions. All experiments presented in this paper were conducted in compliance with the protocol approved by the institutional review board at SUNY College of Optometry and the Declaration of Helsinki, and observers gave written informed consent.

#### Results

Figure 3a shows perceived 3D lengths as a function of 3D pose, averaged over 6 observers and 3 repeats. Dashed lines represent the veridical physical length. Two trends are salient: there is greater underestimation of length for poses pointing towards or away from the observer, and there is greater underestimation of length for longer objects. Individual data show both trends (Figure A2).Underestimation for different object lengths can be compared on the same relative scale in Figure 3b, where the logarithm of the ratio of perceived length over physical length is plotted against the 3D poses. This figure confirms the two trends. The first factor we ruled out for the underestimation is that the dark ground made it ambiguous whether the object was lying on the ground, by rerunning all the conditions of Experiment 1a on a white grid drawn on the ground (Experiment 1b). When the average lengths perceived on the white grid are plotted against average lengths perceived on the dark ground, most symbols fall close to the unit diagonal (Figure 3c), showing that the misestimation of 3D length is very similar on the two ground planes.

**Figure 3.**
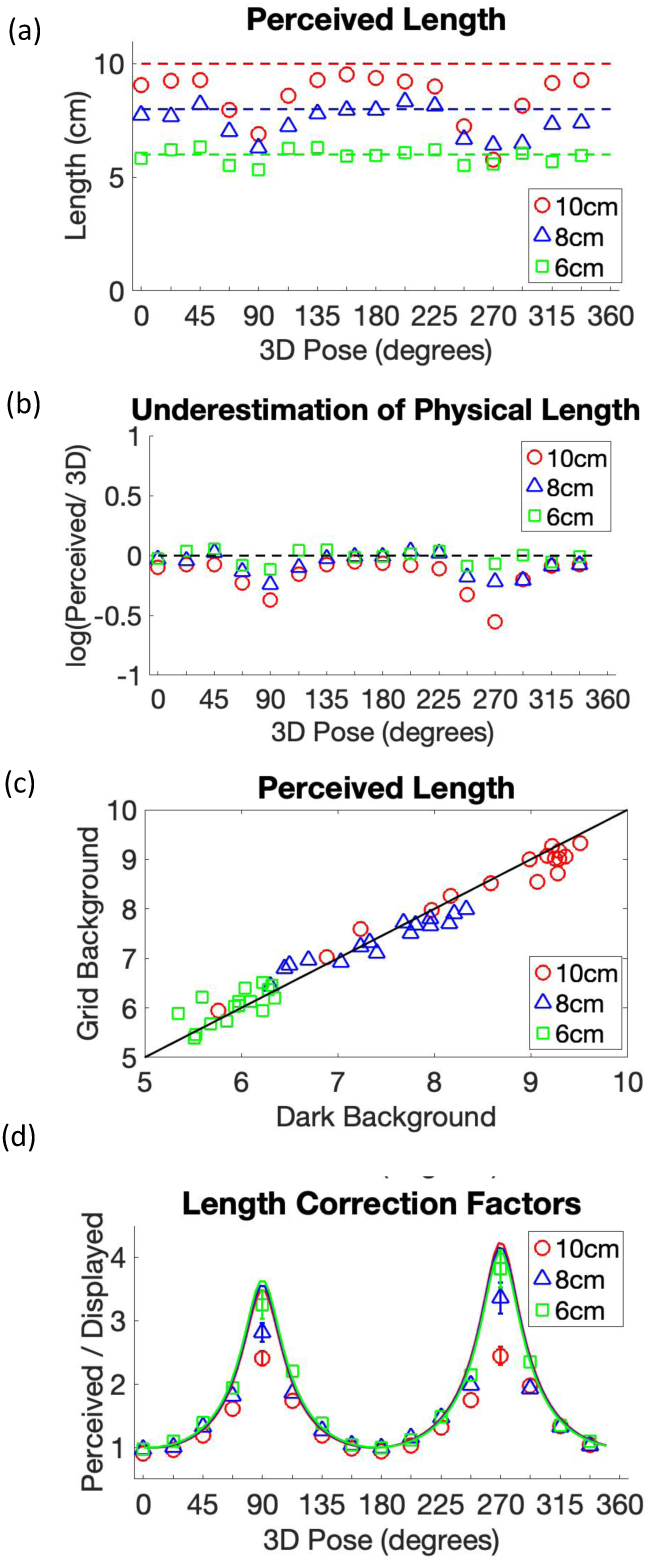
Perceived 3D lengths in Experiment 1. Symbols represent parallelepipeds with lengths 10cm (red circle), 8cm (blue triangle), and 6cm (Green square). (a) Average perceived length across 3D pose (6 observers.) Dashed lines indicate physical 3D lengths. Perceived lengths are underestimated as a systematic function of pose. (b) Logarithm of average perceived 3D length divided by physical length across different 3D poses. Length estimates of the test sticks were close to veridical for front-parallel poses but were seriously underestimated for poses pointing at or away from the viewer (around 90° and 270°.) The underestimation ratio increased with the physical length of the test stick. (c) Perceived length on grid background (Experiment 1b) plotted against perceived length on dark background (Experiment 1a), showing points falling close to the unit diagonal (solid line), indicating that they are similar. (d) Optimal Correction Factor (solid line) and Empirical Correction Factor (symbols) across 3D pose. The empirical correction factor is close to one for the front-parallel poses, which is optimal. The empirical correction factor is greater than one near the line of sight (90° and 270°), but significantly lower than the optimal for the longer sticks. Bars on symbols are 95% confidence intervals.

#### Comparing empirical to optimal size estimation

We now examine underestimation of length for poses towards and away from the observer, especially for longer lengths, by deriving the geometrical information available to observers. For a parallelepiped of length (*L*_3*D*_), the projected length (*L*_*c*_) changes with pose Ω as a distorted sinusoid (viewing elevation = Φ_*c*_, focal length of the camera = *f*_*c*_, and distance from the object = *d*_*c*_):

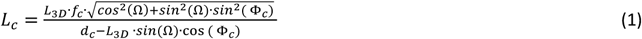

However, the projected length of the vertically oriented cylinder *L*_*m*_ (where the physical length is *L*_3*DM*_), stays invariant with pose because the object is rotated around the axis of the cylinder:

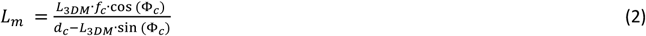

Given the projected lengths on the display, the projected lengths on the retina, *L*_*r*_, would be (focal length of the eye = *f*_*r*_, and distance from the object = *d*_*r*_):

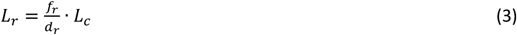

Since the retinal image contains all the information for doing size estimation of these objects, our model predicts 3D size estimates from retinal projections of the objects. Koch et al (2018) showed that humans are excellent at inferring 3D pose of objects on the ground, and their estimations closely match predictions from the geometrical back-projection from retina to the ground plane. In principle, an observer could similarly make veridical estimates of 3D sizes by using the geometrical back-projection from the retinal image, which is given by substituting the expression for *L*_*c*_ from Eq 1 into Eq 3, and then manipulating the equation to get an expression for the optimal estimated 3D length 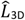:

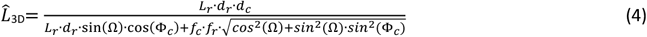

By dividing the optimal length estimate by the projected length, we obtain the Optimal Length Correction index for each pose. The OLC varies little for the three sizes when plotted as a function of 3D pose, as shown by the overlap of the three solid line curves in Figure 3d. The symbols in Figure 3d plot the ratio of perceived length from Figure 3a over projected length (Empirical Length Correction), and show that for poses other than fronto-parallel, observers estimate 3D lengths to be longer than projected lengths (ELC > 1.0). The greatest correction takes place for poses pointing towards or away from the observer, but that is still not sufficient, especially for the longer lengths. The general form of Empirical Length Correction as a function of 3D pose is similar to the Optimal Length Correction curve, suggesting that observers may be using the optimal back-transform, but with an additional multiplicative factor leading to the sub-optimality.

### Geometric model for 3D size estimation

A strong clue to the additional factor is revealed by inspection of the stimuli in Figure 4a. When a 6 cm and a 10 cm parallelepiped are placed on the ground, the upper surface of the longer stick looks more slanted down. The genesis of this illusion is that if the 6 cm stick were increased in length to 10 cm, its projection would come further down along the vertical axis of the image. A similar change in the projected image would happen if the slant of the 6cm stick was increased, or equivalently if the object was pictured from a higher camera elevation. The visual system is thus faced with deciding between an increase in slant versus an increase in length. Figure 4b (left) shows that increasing the camera elevation changes the aspect ratio of a 10cm × 3 cm top surface at 90° pose by a factor of 5.65 from 0.3227 (at 5°) to 1.8247 (at 30°), but barely changes the aspect ratio of the 3cm × 3cm front surface from 1.4120 at 5° to 1.3096 at 30°, a factor of 1.08. The main change is due to the shortening of the projected length of the top surface. The front surface changes are thus not a strong cue to discern change of camera elevation, or equivalently the slant of the object. Thus, it is interesting that visual computations choose to signal a change in slant over a change in length. If a visual system assumes that the longer object is more slanted or pictured from a higher elevation, it will apply a smaller correction factor. Figure 4b (right) shows that the Optimal Length Correction decreases by almost a factor of 5 as the cameral elevation increases from 5° to 30°, thus a misperception of increased slant would lead to a smaller correction factor applied to the projected length.

**Figure 4.**
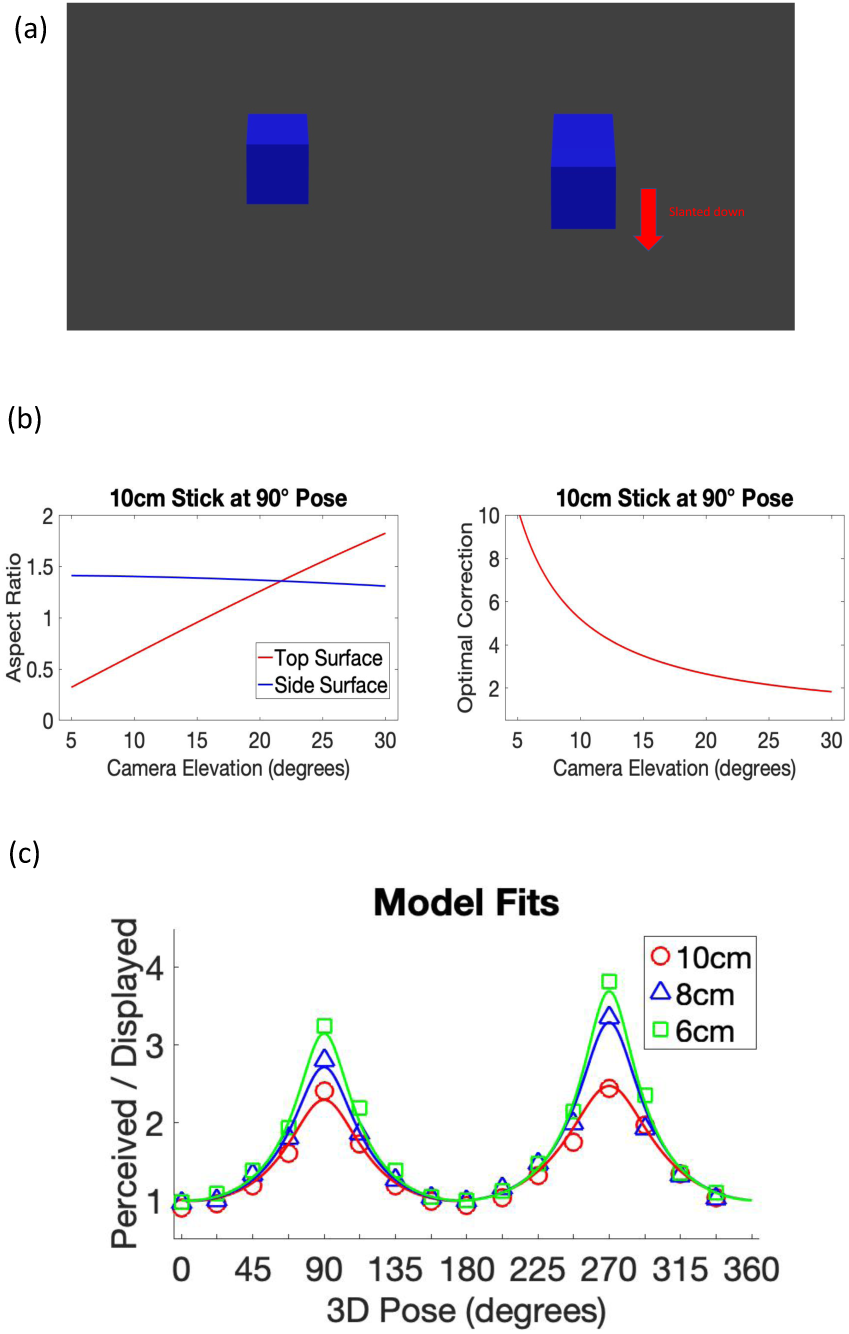
Geometry model for 3D size estimation. (a) 6 and 10 cm test sticks lying on ground, pictured with 15° camera elevation. The longer stick is seen as more slanted down towards the viewer, equivalent to an increase in viewing camera elevation. The effect is the same with a grid on the ground. (b) (left) Aspect ratio of the top surface of 10 × 3 cm (red) and front surface of 3×3 cm (blue) of stick lying on ground in 90 degree pose against camera elevation. The main change is the projection of the length of the top surface. (right) Optimal Correction Factor for the stick as a function of camera elevation. Correction factor decreases with increasing camera elevation. (c) A model using the optimal geometrical back-transform but with overestimation of viewing elevation fits the underestimates of object length.

Based on the results and the visual observations, we formulated the hypothesis that observers are using an optimal back-transform, but overestimating the slant of the object (or equivalently the camera elevation), thus correcting less than required for the shortening. Therefore, we tested whether adding a multiplier *k*>1 to the viewing elevation in the optimal geometrical back-transform function could provide good fits to the empirical estimates for different physical lengths:

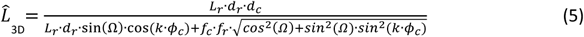

Figure 4c replots the empirical corrections from Figure 3d for the three lengths of test sticks. For each length, we found the *k* for which the optimal correction factor curve best fits the empirical correction factors, and shows that the model fits the results for all three physical lengths well, with just one free parameter that increases perceived camera elevation. Since perceived slant can be slightly different for 90° and 270° poses, we allowed *k* to be different values for 0-180 and 180-360. Based on the best fitting *k* values, the estimated camera elevations were around 16° for the 6 cm stick, 20 for the 8 cm, and 25 for the 10 cm. Parenthetically, we also tested whether putting a multiplier on distance, to simulate misperceived distance (Sedgwick, 1989), could explain the data, but that was not successful. Consequently, we tested whether the slant illusion would hold up to quantitative measurements.

### Experiment 2: Slant misestimation as factor for suboptimal length correction

There is no way to measure absolute perceived slants with good precision, so we checked whether relative perceived slants across stick lengths followed the trend predicted by the best estimated camera elevations in the model. We compared fixed perceived slants of 6, 8 and 10 cm sticks to the adjustable perceived slant of a 6 cm stick, when both sticks were either at 90° poses or at 270° (Figure 5a).

**Figure 5.**
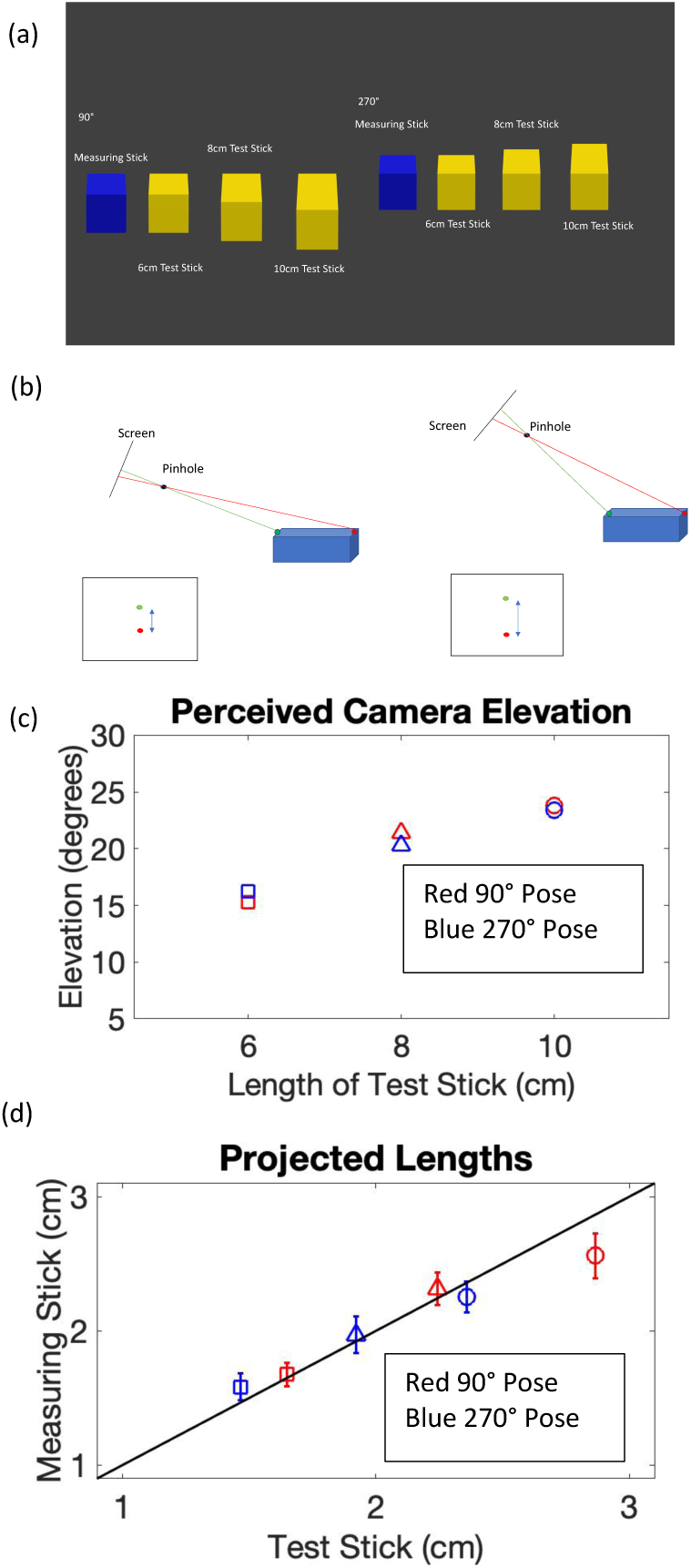
Slant misestimation as a function of length. Experiment 2: (a) Blue adjustable measuring stick (6cm) was paired with one yellow test stick (10, 8, or 6cm) lying on dark ground in in the same pose (90° or 270°). (b) Observers were asked to match slant, but they were actually changing camera elevation on just the measuring stick (Note change in angle between camera screen and top surface of object). (c) Relative perceived camera elevation for each length of fixed stick (circle-10cm, triangle-8cm, square-6cm), separately for the two poses (red-90°, blue-270°). Observers’ overestimation of camera elevation increases with the length of test stick, which corresponds to an increase in perceived slant. (d) At the slant match, projected lengths (vertical extents in image) for the two sticks are roughly equal (bars indicate 95% confidence intervals).

#### Methods

The observers from Experiment 1 also participated in Experiment 2. Two kinds of images were made by the Blender. The yellow fixed-stick’s image was rendered from a 15° camera elevation. A series of images were rendered for the blue adjustable-stick, from 0° to 30° elevation at every 1° step. The other properties of the Camera were the same as Experiment 1, as were the display monitor and observer viewpoint settings. Observers were asked to adjust the slant of the blue stick to match that of the paired yellow fixed stick by pushing buttons, without time limits. Unknown to them, observers were actually adjusting the camera elevation on the rendering of the blue stick through the 31 possible settings. Figure 5b shows that this manipulation changes the angle between camera screen and top surface of the object, so it has the same effect as changing the slant of the object. Since the aspect ratio of the front surface barely changes in these settings (Figure 4b left), the front surface provides almost no clue to the relative slant for these adjustments. The two sticks were randomly assigned to left and right on each trial. Three separate sets contained random assignments of every condition (2 poses of the pair × 3 lengths of fixed stick). Observers were allowed to take a break between sets.

#### Results

The main result (Figure 5c) is that observers perceived camera elevations as higher for the longer fixed sticks, despite all sticks lying flat on the same ground plane. When both sticks were posed at 90°, the average camera elevations were and 15.278°, 21.39° and 23.83° for the 6 cm, 8 cm, and 10 cm sticks respectively. For the 270° poses, the averages were 16.22°, 20.33° and 23.39° respectively. Individual results are shown in Figure A3. These values are not exactly the same as we estimated for the best fits of the model, because we measured relative slants and the model incorporates absolute perceived slants. However, the results are compatible with our hypothesis that observers may be applying a smaller correction factor to longer sticks because they see them as more slanted. It is interesting that when equating slant, observers end up also equating projected lengths of the two top surfaces, i.e. the excursion along the vertical axis of the image (Figure 5d).

That the percept of increased slant is very compelling is demonstrated by the movie in Figure 6. We began with the same rotating object as in Figure 1, but used the fitted model in Figure 4c to dynamically adjust the lengths of the limbs so they are perceptually equal in all poses. As the object rotates, the limb facing towards or away from the observer appears to bounce up and down because the slant illusion seems to dominate the percept of changing length. In the dynamic case, this could be related to the possibility that articulated objects are more common in the world then are objects that increase or decrease in length over a short period. The same appeal to natural statistics could be invoked to explain the difficulty in perceiving expansion or contraction in depth of solid objects (Johansson, 1964; Jansson & Johansson, 1973; Jain & Zaidi, 2011), while other deformations are easy to discern, even for rotating and flowing shapes (Cohen, Jain & Zaidi, 2010; Fantoni, Caudek & Domini, 2014; Bates et al, 2019). However, the slant illusion is just as compelling in the static case, so biases in perceiving depth versus extent from retinal images may also be in play (Jain & Zaidi, 2013; Kim & Burge 2018). Although not a part of this study, we want to note that perceived slant is also affected by other factors, such as object shape. For example, 6, 8 and 10 cm cubes are much more similar in apparent slant than the three sizes of parallelepipeds used in this study, and informal manipulations suggest that length to height ratio is also a factor.

**Figure 6.**
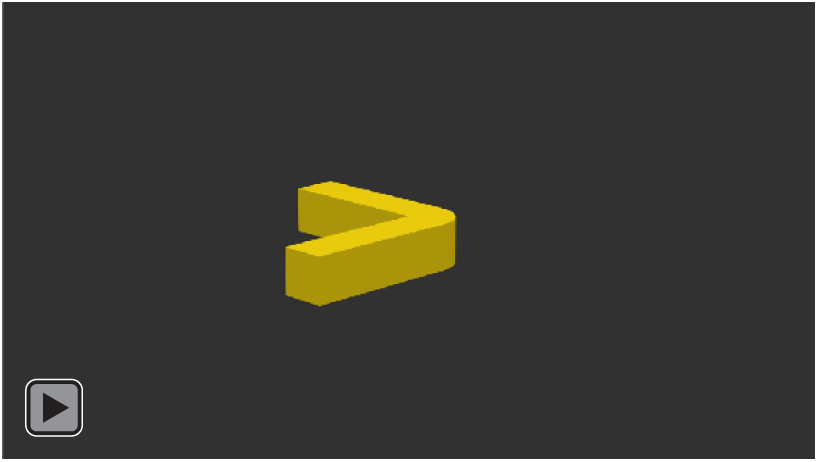
Demonstration of slant illusion. A rigid object with two perceptually equal arms at a right angle is rotating on the ground. Each arm seems to bounce up and down when it faces towards or away from the observer, because the slant illusion dominates percepts of length changes.

## Discussion

The first empirical contribution of this study is to measure size estimates of 3D objects as they are rotated on a ground plane into different poses, which is equivalent to changing viewpoints around the object. Invariance to rotation is one of the mathematical properties of shape, so it is surprising that no previous studies looked at constancy or inconstancy of shape or relative size across poses. We found systematic and repeatable distortions of perceived size across poses even for simple symmetric and regular objects. Estimates of length were close to veridical for fronto-parallel poses but were seriously underestimated for poses pointing at or away from the observer. Interestingly, the underestimation increased with the physical length of the parallelepiped. Slant matching measurements revealed that longer objects were seen as slanted down, equivalent to an increase in viewing elevation. The second empirical contribution of the study is this slant illusion, and the demonstration that the visual system seems to prefer an illusory slant to an actual length change, when slanting or lengthening an object lead to very similar images.

The main theoretical contribution of this study is to link 3D size estimates to the mental use of projective geometry. Size estimates as a function of pose follow a similar shaped curve as the optimal back-transform would predict. Thus, our model that incorporates observers’ misestimates of object slant in the optimal geometrical back-transform equation can explain the inconstancy of relative size for different poses, and for different object sizes. Thus, size inconstancy results despite observers using the correct geometric back-transform, if retinal images evoke misestimates of slant. Remarkably, our observers made similar relative size estimates across different poses, suggesting that the mental sub-conscious use of this geometrical knowledge is common among all observers.

Although we have expressed our measurements in terms of length, our measurements also reveal inconstancy of shape perception across poses, or equivalently different views of one static object. Consider the dual-limbed object formed by the parallelepiped and cylinder together in Figure 2. Since the perceived length of the parallelepiped changes in different poses relative to the perceived length of the cylinder, the object is not perceived as constant in shape. Another example of perceived shape changes, because of relative limb lengths not being constant, is provided by the movie in Figure 1. In addition, our measurements predict that a solid 3-D object will undergo perceived aspect ratio changes in different poses, as perceived size along the axis pointing towards the observer will be underestimated compared to the two orthogonal axes. Our results suggest that observers do try to correct for the projected shortening, using knowledge of projective geometry, but it is not enough to achieve shape constancy. The inconstancy of estimated depth relative to width, found by previous studies on perceived object shape and perceived scene distances, may also be explained by our geometrical back-transform model.

It is theoretically interesting that we show that humans do not completely discount the distortions created by perspective projection, despite possibly using the optimal geometric back transform. It is worth considering what else could be explained by assuming mental knowledge of projective geometry. Projective geometry preserves continuity, collinearity and convergence. If a visual system assumes that when collinear edges and intersections between edges occur in an image, it is generally because the perspective projection is preserving continuous edges and corners from the 3D world, then some objects separated into random parts, e.g. Ames Chair (Ittelson, 1952), would be seen as unbroken and cohesive from the proper viewpoint, without having to invoke stronger assumptions of simplicity or regularity in reconstructing the 3D world (Li & Pizlo, 2011). The “generic viewpoint assumption” (Freeman, 1994) thus implicitly contains a “generic projective geometry assumption”, and this can be made more concrete in object and scene perception by incorporating priors that assume the invariants of projective geometry, especially when extracting 3D shapes from contours (Li, Pizlo & Steinman, 2009; Elder, 2018; Wang et al, 2018). The fact that slant is ambiguous in perspective projection is compatible with some real-world illusions of slant and non-rigidity (Griffiths & Zaidi, 1998, 2000). Animals and humans have constant exposure to perspective projection through image-forming eyes. Therefore, it has been an open question whether brains have learned to exploit projective geometry to understand 3D scenes. Our results imply that human brains use embedded knowledge of projective geometry to estimate 3D sizes and shapes from their retinal images. This adds to our previous results that humans use optimal projective geometry back-transforms to estimate 3D poses in real 3-D scenes and their pictures (Koch et al 2018). Humans thus seem to have internalized particular aspects of projective geometry through evolution or learning.

## Appendix

### Calibrating sizes rendered by Blender

Theoretical projected lengths calculated from Eqs 1 and 2 were compared with the lengths of the sticks rendered by Blender on the display screen. Since our main concern in Experiment 1 was with the relative lengths of the horizontal and vertical sticks in the 3D scenes, we calculated the ratios of the projected lengths of the parallelepiped to the orthogonally attached cylinders of the same physical length and plotted them as a function of 3D pose. Blender and geometrically derived ratios both followed a distorted and asymmetric sinusoidal curve, and had very similar values (Figure A1). Since the derived projection of the measuring stick is always the same length, this curve also describes the projected length of the test stick as a function of pose.

### Lengths estimated by individual observers

Individual observer’s length estimates corresponding to Figure 3a are shown in Figure A2.

### Misestimates of slant by individual observers

Individual observer relative slant estimates corresponding to Figure 5c are shown in Figure A3.

## ACKNOWLEDGMENTS

This work was supported by National Institutes of Health Grants EY13312 and EY07556. We are grateful to Erin Koch for many discussions about all aspects of this study.

## Figure legends (Appendix)

**Figure A1.**
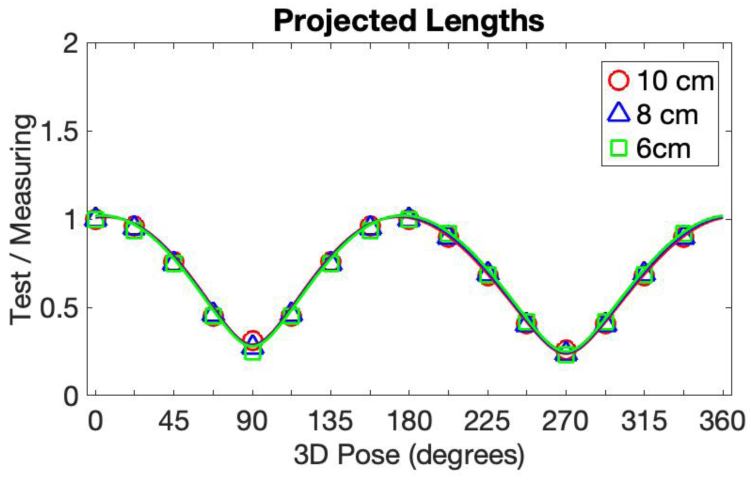
Calibrating rendered images for relative size. Projected length of Test Stick divided by projected length of an equal sized Measuring Stick. There is no difference in the derived ratios for the three physical lengths shown as solid line curves. Points represent physical measurements of lengths of objects rendered by Blender on display screen.

**Figure A2.**
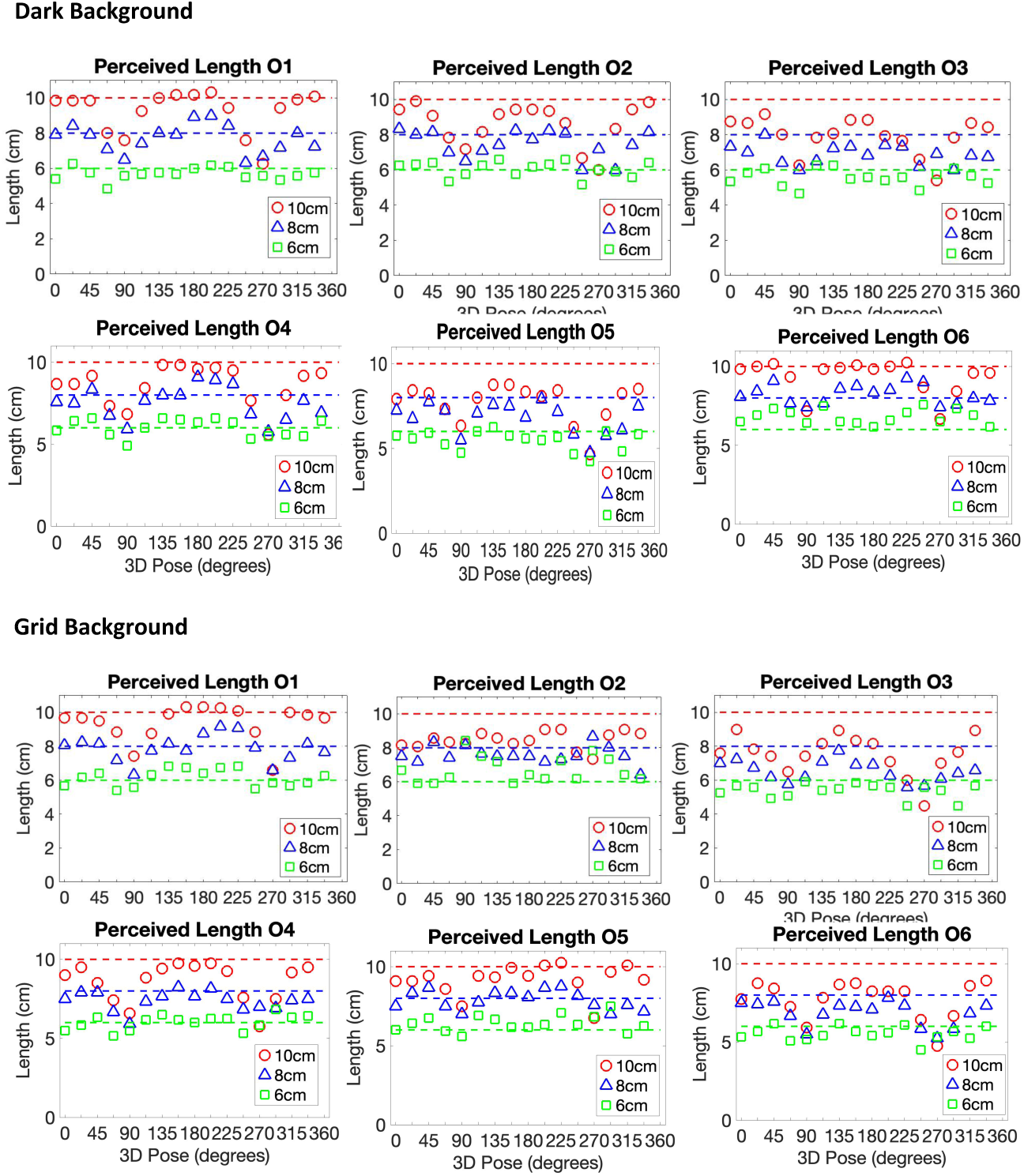
Perceived 3D length in Experiment 1 for each observer. Symbols represent parallelepipeds with lengths 10cm (red circle), 8cm (blue triangle), and 6cm (Green square). Average perceived length across 3D pose (6 observers.) Dashed lines indicate physical 3D lengths. Perceived lengths are underestimated as a systematic function of pose. The systematic patterns are similar for all observers.

**Figure A3.**
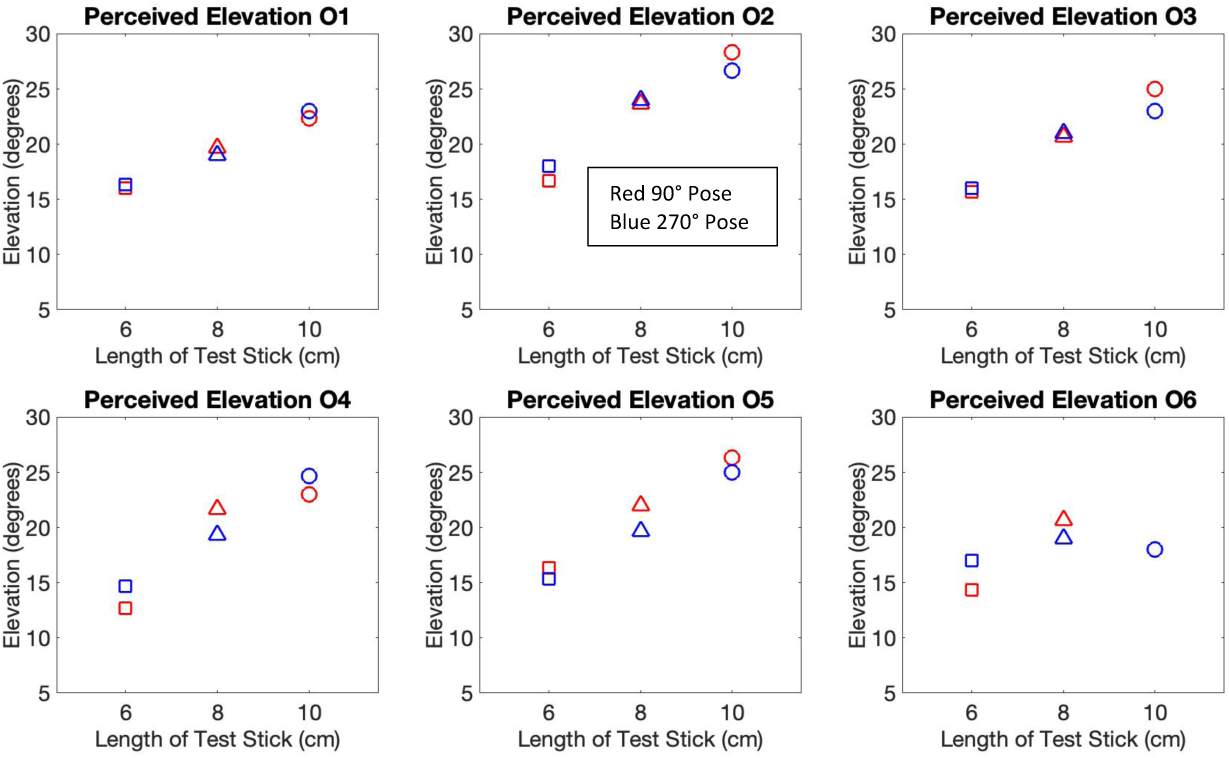
Slant misestimation as a function of length for each observer. Blue adjustable measuring stick (6cm) was paired with one yellow test stick (10, 8, or 6cm) lying on dark ground in the same pose (90° or 270°). Relative perceived camera elevation for each length of fixed stick (circle-10cm, triangle-8cm, square-6cm), separately for the two poses (red-90°, blue-270°). Observers’ overestimation of camera elevation increases with the length of test stick, which corresponds to an increase in perceived slant.

